# Yeast heterochromatin regulators Sir2 and Sir3 act directly at euchromatic DNA replication origins

**DOI:** 10.1101/271304

**Authors:** Timothy A Hoggard, FuJung Chang, Kelsey Rae Perry, Sandya Subramanian, Jessica Kenworthy, Julie Chueng, Erika Shor, Michael Cosgrove, Jef D. Boeke, Catherine A Fox, Michael Weinreich

## Abstract

Most active DNA replication origins are found within euchromatin, while origins within heterochromatin are often inactive or inhibited. In yeast, origin activity within heterochromatin is negatively controlled by the histone H4K16 deacetylase, Sir2, and at some heterochromatic loci also by the nucleosome binding protein, Sir3. The prevailing view has been that direct functions of Sir2 and Sir3 are confined to heterochromatin. However, growth defects in yeast mutants compromised for loading the MCM helicase, such as *cdc6-4*, are robustly suppressed by deletion of either *SIR2* or *SIR3*. While this and other observations indicate that *SIR2,3* can have a negative impact on at least some euchromatic origins, the genomic scale of this effect was unknown. It was also unknown whether this suppression resulted from direct functions of Sir2,3 within euchromatin, or was an indirect effect of their previously established roles within heterochromatin. Using both MCM ChIP-Seq and MNase-H4K16ac ChIP-Seq data, we show that a *SIR2* deletion rescues MCM complex loading at ~80% of euchromatic origins in *cdc6-4* cells. Therefore, Sir2 exhibits a pervasive effect at the majority of euchromatic origins. Importantly, in wild type (i.e. *CDC6*) cells, origin-adjacent nucleosomes were depleted for H4K16 acetylation in a *SIR2*-dependent manner. In addition, both Sir2 and Sir3 directly bound to nucleosomes adjacent to euchromatic origins. The relative levels of each of these molecular hallmarks of yeast heterochromatin – *SIR2*-dependent H4K16 hypoacetylation, Sir2, and Sir3 – correlated with how strongly a *SIR2* deletion suppressed the MCM loading defect in *cdc6-4* cells. Finally, a screen for histone H3 and H4 mutants that could suppress the *cdc6-4* growth defect identified amino acids that map to a surface of the nucleosome important for Sir3 binding. We conclude that heterochromatin proteins directly bind euchromatic DNA replication origins and modify their local chromatin environment.

## AUTHOR SUMMARY

When a cell divides, it must copy or “replicate” its DNA. DNA replication starts at chromosomal regions called origins when a collection of replication proteins gains local access to unwind the two strands. Chromosomal DNA is packaged into a protein-DNA complex called chromatin and there are two major structurally and functionally distinct types. Euchromatin allows DNA replication proteins to access origin DNA, while heterochromatin inhibits their access. The prevalent view has been that the heterochromatin proteins required to inhibit origins are confined to heterochromatin. In this study, the conserved heterochromatin proteins, Sir2 and Sir3, are shown to both physically and functionally associate with the majority of origins in euchromatin. This raises important new questions about the chromosomal targets of heterochromatin proteins, and how and why the majority of origins exist within a potentially repressive chromatin structure.

## Introduction

In eukaryotic cells, efficient genome duplication requires the function of multiple DNA replication origins distributed over the length of each chromosome [1–5]. Origins are selected by a series of steps in late M- to G1-phase during which the origin recognition complex (ORC) binds directly to DNA and recruits the Cdc6 protein [6–8]. The ORC- Cdc6-DNA complex recruits Cdt1-MCM to form an MCM double hexamer (dhMCM) encircling double-stranded DNA [9,10]. Several kinases and loading factors then remodel the dhMCM into two active CMG helicases (<u>C</u>dc45<u>-M</u>CM<u>-G</u>INS) that unwind the DNA bidirectionally from each origin to allow the initiation of DNA synthesis [11]. Thus, dhMCM loading is the event that ‘licenses’ the DNA to function as an origin of replication in S-phase (for recent comprehensive review of yeast replication see [12]). All replication origins exist in the context of chromatin, and origin function is significantly affected by local chromatin structure (reviewed in [13,14]). Typically, the most efficient origins (i.e. the origins that are used in most cell cycles) are found in euchromatin, while less efficient origins are associated with heterochromatin. While there is intense interest in the impact of chromatin structure on origin function, the relevant molecular features of origin-adjacent chromatin and the steps in origin function that they control remain incompletely understood.

Yeast heterochromatin is characterized by hypoacetylated nucleosomes and associated heterochromatin regulatory proteins that promote a compact chromatin structure that makes the underlying DNA inaccessible to protein-DNA interactions and processes such as transcription and DNA replication initiation (recently reviewed in [15]). The Sir2 (**S**ilent **i**nformation **r**egulator) protein, the founding member of a family of conserved NAD-dependent protein deacetylases, removes an acetyl group from lysine 16 of histone H4 (H4K16) and is required for heterochromatin formation in budding yeast [16–19]. Heterochromatin domains form at only a few discrete regions in the yeast genome, rDNA, telomeres, and the *HM*-mating type loci, and at each of these loci, Sir2 both deacetylates H4K16ac and stably binds to nucleosomes [15]. Heterochromatin formation at the *HM* loci and telomeres also requires nucleosome binding by the Sir3 and Sir4 proteins.

While it is recognized that the function of normally efficient origins can be suppressed when they are engineered within heterochromatic regions of the genome [20,21], several more recent studies reveal that *SIR2* can also affect the function of euchromatic origins [22,23]. However, depending on experimental context, *SIR2* can be interpreted to exert either positive or negative effects on euchromatic origins. In particular, recent studies reveal that *SIR2* acts indirectly as a positive regulator of euchromatic origins by suppressing the function of rDNA origins [24,25]. The tandemly repeated rDNA gene locus on chromosome XII contains ~200 replication origins, but ~80% of these origins are suppressed by Sir2-dependent heterochromatin [24]. Because the rDNA origins account for >30% of all yeast genomic origins, deletion of *SIR2* results in activation of many rDNA origins, which then sequester limiting S-phase origin activation factors from euchromatic origins, delaying their activation time in S-phase [26,27]. Thus, the positive role for *SIR2* in regulating euchromatic origin function in these reports is explained as a byproduct of a direct inhibition of rDNA origins by Sir2.

Interestingly, other studies establish that *SIR2* can also exert a negative effect at euchromatic origins. This negative role was revealed by a classic genetic screen that isolated *sir2* mutants as suppressors of the temperature-sensitive *cdc6-4* allele [23,28,29]. Cdc6, a member of the AAA+ protein family, must bind ATP to load dhMCM [30,31]. The *cdc6-4* allele encodes a mutant Cdc6 with a lysine to alanine substitution in the conserved ATP binding motif. Cells with the *cdc6-4* allele are viable but have S-phase associated growth defects at the permissive temperature and arrest growth at the non-permissive temperature with a failure to load dhMCM, as assessed by origin-specific ChIPs. Like *sir2Δ*, *sir3Δ* is also a robust suppressor of the *cdc6-4* growth defect, whereas a *sir4Δ* mutant is only a very weak suppressor [29]. Thus the suppression by *sir2Δ* or *sir3Δ* is not easily explained by defects in classic Sir-heterochromatic gene silencing of known loci because Sir2, Sir3 and Sir4 are each equally essential for *HM*- and telomere silencing and only Sir2 is required for rDNA silencing. In addition, multiple *sir2* alleles specifically defective in rDNA silencing do not suppress *cdc6-4* ts lethality, indicating that a loss of rDNA silencing is not sufficient to explain *SIR2’s* negative effect on euchromatic origins [29]. These findings support a distinct, rDNA- and classic heterochromatin-independent function of *SIR2* on euchromatic origins. Importantly, a *sir2* catalytic mutant or a histone mutant that converts H4K16 into a residue that mimics acetylated H4K16 (H4K16Q) suppresses *cdc6-4*, indicating that the Sir2 deacetylase function is critical for its negative impact on euchromatic origins [23,29]. However, these data do not address whether the negative role exerted by *SIR2* is due to direct roles for Sir2 within euchromatic regions of the genome, or whether *SIR2* and *SIR3* affect euchromatic origins through a shared mechanism.

In this report we investigated the molecular mechanisms by which *SIR2* affected euchromatic origins by performing MCM ChIP-Seq and MNase-H4K16ac ChIP-Seq experiments and also by analyzing several published high-resolution ChIP-Seq experiments [32–34]. Our results provide evidence for a direct role for Sir2 and Sir3 in forming a repressive local chromatin environment around most origins that exist within euchromatin. A *sir2Δ* mutation was sufficient to restore MCM binding at the majority of euchromatic origins in *cdc6-4* cells even at the non-permissive growth temperature for this allele. In wild type cells, nucleosomes immediately adjacent to the majority of euchromatic origins were relatively hypoacetylated on H4K16 compared to non-origin control loci, a behavior unique to this histone acetylation mark. Moreover, this origin-specific H4K16 hypoacetylation was completely dependent on *SIR2*. In addition, Sir2 and Sir3 were physically associated with euchromatic origins but not non-origin control loci. The levels of these three distinct molecular features of yeast heterochromatin—hypoacetylation of H4K16, Sir2 and Sir3 binding—correlated with how strongly a *SIR2* deletion suppressed the MCM loading defect in *cdc6-4* cells. Based on these results we propose that the dhMCM loading reaction has evolved to work within a potentially repressive Sir2-Sir3 chromatin environment that forms around most euchromatic origins. Consistent with this model, a screen for suppressors of *cdc6-4* temperature-sensitive lethality identified several histone mutants known to disrupt the Sir3-nucleosome binding interface.

## Results

***SIR2* inhibited MCM loading at most euchromatic origins in *cdc6-4* mutant cells**. To examine the extent to which a *SIR2* deletion (*sir2Δ*) suppresses defects in MCM loading caused by the *cdc6-4* allele, we assessed MCM binding by ChIP-Seq in four congenic yeast strains: wild type (*SIR2 CDC6*); *sir2Δ*; *cdc6-4*; and *sir2Δ cdc6-4*. Because *sir2Δ* rescues the temperature-sensitive growth defect of *cdc6-4* [29], the experiment was performed using chromatin from cells incubated at 37°C, the non-permissive growth temperature for *cdc6-4* (**Figure 1A**). Cells were arrested in M-phase at the permissive growth temperature (25°C), shifted to 37°C, and then released into G1-phase to allow time for MCM loading. MCM ChIP-Seq was performed for each strain using a monoclonal antibody against Mcm2 [23]. Examination of MCM ChIP-Seq signals across the genome, as illustrated for chromosomes III and VI, revealed that MCM distribution in wild type and *sir2Δ* cells was very similar (**Figure 1B, 1C**, **S1 Figure**). *Cdc6-4* cells failed to produce MCM ChIP-Seq signals above background levels, consistent with the essential role of Cdc6 in MCM loading **(Figure 1B, 1C**). In contrast, *cdc6-4 sir2Δ* mutant cells showed MCM association, albeit to varying degrees, at most origins. (**Figure 1B, C**, **S1 Figure**). Therefore, deletion of *SIR2* rescued origin-specific association of MCM at the majority of chromosomal origins in *cdc6-4* mutant yeast, consistent with *SIR2* have a pervasive negative effect at the majority of euchromatic origins.

A previous study screened for origins on chromosomes III and VI (**Figure 1**, gray boxes) that, when cloned onto a plasmid, were functional in *cdc6-4 sir2Δ* cells growing at the non-permissive temperature for *cdc6-4*. Five origins were identified as *SIR2*- responsive in this screen: *ARS317*, the *HMR*-E silencer origin, and four additional origins *ARS305*, *ARS315*, *ARS603* and *ARS606* (**Figure 1B, C** **and D**, black boxes). *ARS317* was not used in subsequent analyses here because it did not produce a robust MCM ChIP-Seq signal in wild type cells, and because we eliminated all origins that exist within heterochromatic domains for deeper analyses, as described below. The plasmid-based study suggested that at least 20% of yeast origins were likely to be *SIR2*-responsive [23]. In contrast, the MCM-ChIP-Seq revealed that ~80% of origins showed MCM binding in *cdc6-4 sir2Δ* cells (**Figure 1B, 1C**, **and S1 Figure**). We note that the plasmid-based assay demanded that origin function was rescued to a level in *cdc6-4 sir2Δ* cells that allowed for colonies to form at 37°C, a selection that was likely more stringent than the screen for an MCM ChIP-Seq signal as in Figure 1. In this regard, it is notable that the origins identified as *SIR2*-responsive in the plasmid screen were indeed among those that showed the most robust rescue of MCM ChIP-Seq signals in *cdc6-4 sir2Δ* cells (**Figure 1D**, black boxes). In addition, the plasmid-based screen assessed only small origin fragments that might not have recapitulated a nucleosome organization required to show *SIR2*-responsiveness. Indeed, plasmid-born *ARS1005* (a top origin identified by MCM ChIP-Seq) exhibited *SIR2*-regulation only when a larger chromosomal region surrounding *ARS1005* was present that could accommodate the adjacent chromosomally directed nucleosomes (**S2 Figure**).

**Figure 1.**
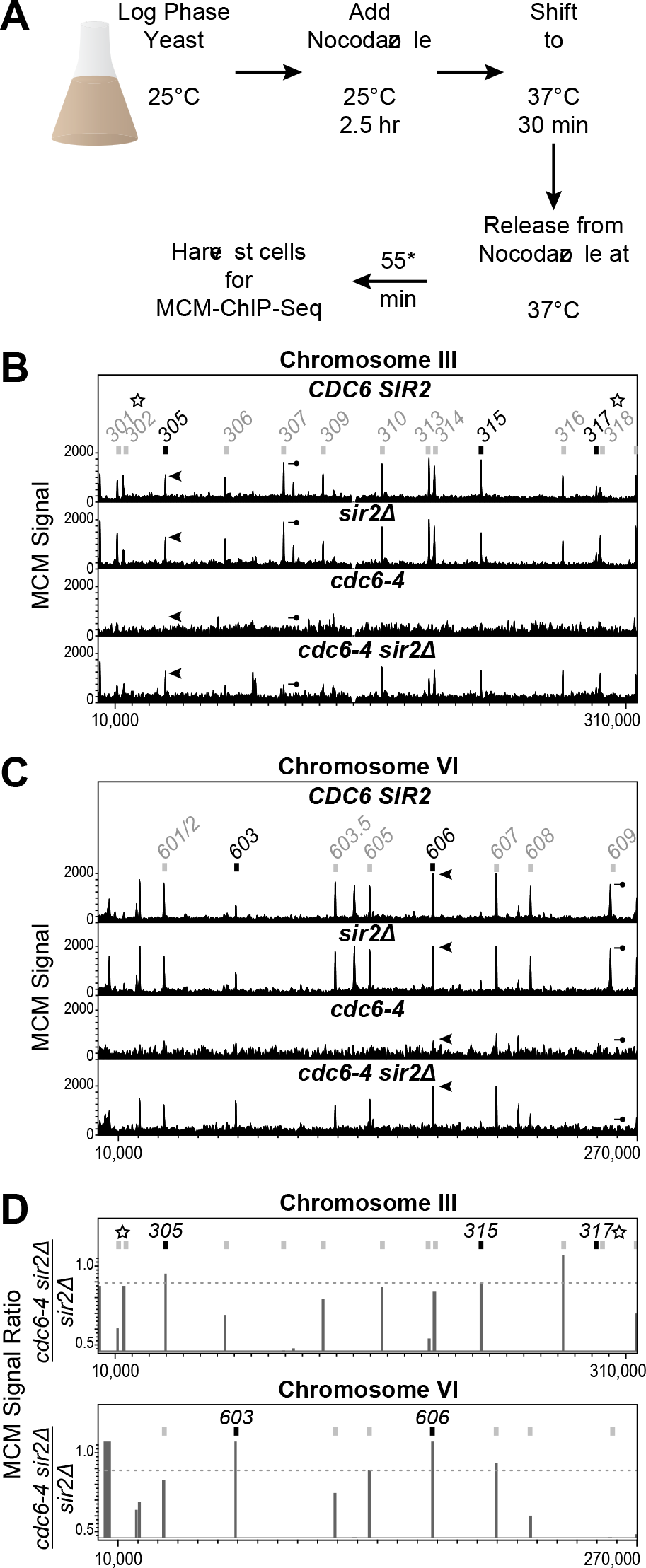
*SIR2* inhibited MCM association at the majority of euchromatic origins in *cdc6-4* mutant cells. **A.** Experimental outline: MCM binding to chromosomal DNA was examined by ChIP-Seq from congenic *CDC6 SIR2* (wild type), *CDC6 sir2Δ* (*sir2Δ*), *cdc6-4 SIR2* (*cdc6-4*), and *cdc6-4 sir2Δ* cells. Cells were released from a G2/M nocodazole-arrest into G1 phase at 37oC prior to formaldehyde crosslinking. **B.** MCM ChIP-Seq signals across Chromosome III. The x-axis indicates the chromosomal coordinates, the y-axis the normalized MCM read counts (MCM signal). The scale on the y-axis is the same for each of the four strains. Confirmed origins on Chromosome III are indicated by boxes and numbers. Black boxes indicate origins that were defined as *SIR2*-responsive because they provided for plasmid replication in *cdc6-4 sir2Δ* cells at the non-permissive temperature for *cdc6-4* [23]. The arrowhead and the line-with-circle indicate *ARS305* and *ARS307*, respectively that produced similar association with MCM in wild type cells but different degrees of MCM association in *cdc6-4 sir2Δ* cells. These origins are highlighted to illustrate that the wild type pattern of MCM origin distribution was not rescued fully in *cdc6-4 sir2Δ* cells. The starred origins (*ARS301*, *ARS302*, *ARS317* and *ARS318*) are associated with transcriptional silencers that direct the assembly of SIR-heterochromatin at the *HML* and *HMR* loci on chromosome III [15]. They were excluded from further analyses in this study for this reason, as discussed in text. **C.** MCM ChIP-Seq signals across Chromosome VI. The arrowhead and the line-with-circle indicate *ARS606* and *ARS609*, respectively, which behaved analogously to *ARS305* and *ARS307*, respectively, as described in B. **D.** Origins’ *SIR2*-responsiveness was defined as the ratio of the MCM signal at the origin in *cdc6-4 sir2Δ* cells relative to *sir2Δ* cells. We confined analyses to origins to that produced strong signals in both WT and *sir2Δ* cells as defined by inclusion among the top 400 chromosomal coordinates of enrichment (**S1 Figure**). This approach successfully identified the euchromatic origins (*ARS305*, *ARS315*, *ARS603 and* ARS606) that had previously been defined as *SIR2*-responsive origins based on a systematic plasmid-based screen of these chromosomal origins [23].

**Reduction in rDNA copy number did not explain suppression of *cdc6-4* by *sir* mutants**. In yeast, each rDNA repeat contains a single rDNA origin, and these repeats are present in hundreds of tandem copies per cell on chromosome XII (**Figure 2A**). Some mutants that affect origin function, such as *orc2-1*, can be suppressed by reducing the rDNA origin-load [35,36]. Therefore, we tested whether reduced rDNA copy number might account for suppression of *cdc6-4* by *sir2Δ* by analyzing the rDNA locus in the strains used for the MCM ChIP-Seq using qPCR (**Figure 2B**). While rDNA copy number varied from colony to colony even within a single strain, on average, wild type, *sir2Δ* and *cdc6-4* cells had similar levels of rDNA (**Figure 2C**). However, *cdc6-4 sir2Δ* cells showed a 2-fold reduction in rDNA levels relative to *cdc6-4* cells (P = 0.02), raising the possibility that suppression of *cdc6-4* by *sir2Δ* was mediated at least in part by reductions in rDNA copy number (**Figure 2C**). To further address this issue, we performed two additional experiments. First, we exploited the previous observation that *sir3Δ* also suppresses *cdc6-4* [29]. In contrast to *SIR2*, *SIR3* has no role in rDNA silencing [37,38]. *Cdc6-4 sir3Δ* cells grew at the non-permissive temperature for *cdc6-4*, as expected [29] (**Figure 2D**). However, the rDNA copy number of *cdc6-4 sir3Δ* yeast was slightly higher compared to *cdc6-4* yeast, indicating that reduced rDNA copy number could not explain this suppression (**Figure 2E**). Second, we asked whether a reduction in rDNA copy number was sufficient to suppress *cdc6-4* by generating *cdc6-4* cells with only 35 copies of rDNA (rDNA-35) (**Figure 2D**). These cells also contained a *FOB1* deletion to reduce amplification of the rDNA locus [39,40]. The *cdc6-4 rDNA-35 fob1Δ* cells failed to grow at the non-permissive temperature and grew no better than the *cdc6-4 rDNA-180 fob1Δ* cells (**Figure 2D**). Thus, reducing rDNA copy number was neither necessary nor sufficient to explain *SIR*-mediated suppression of *cdc6-4*. Cells harboring a reduction in rDNA copy number in *cdc6-4 sir2Δ* populations likely arise because loss of Sir2 increases recombination frequency within the rDNA array, and rDNA copy numbers are reduced over time in many mutants compromised for replication initiation [35,41,42].

**Figure 2.**
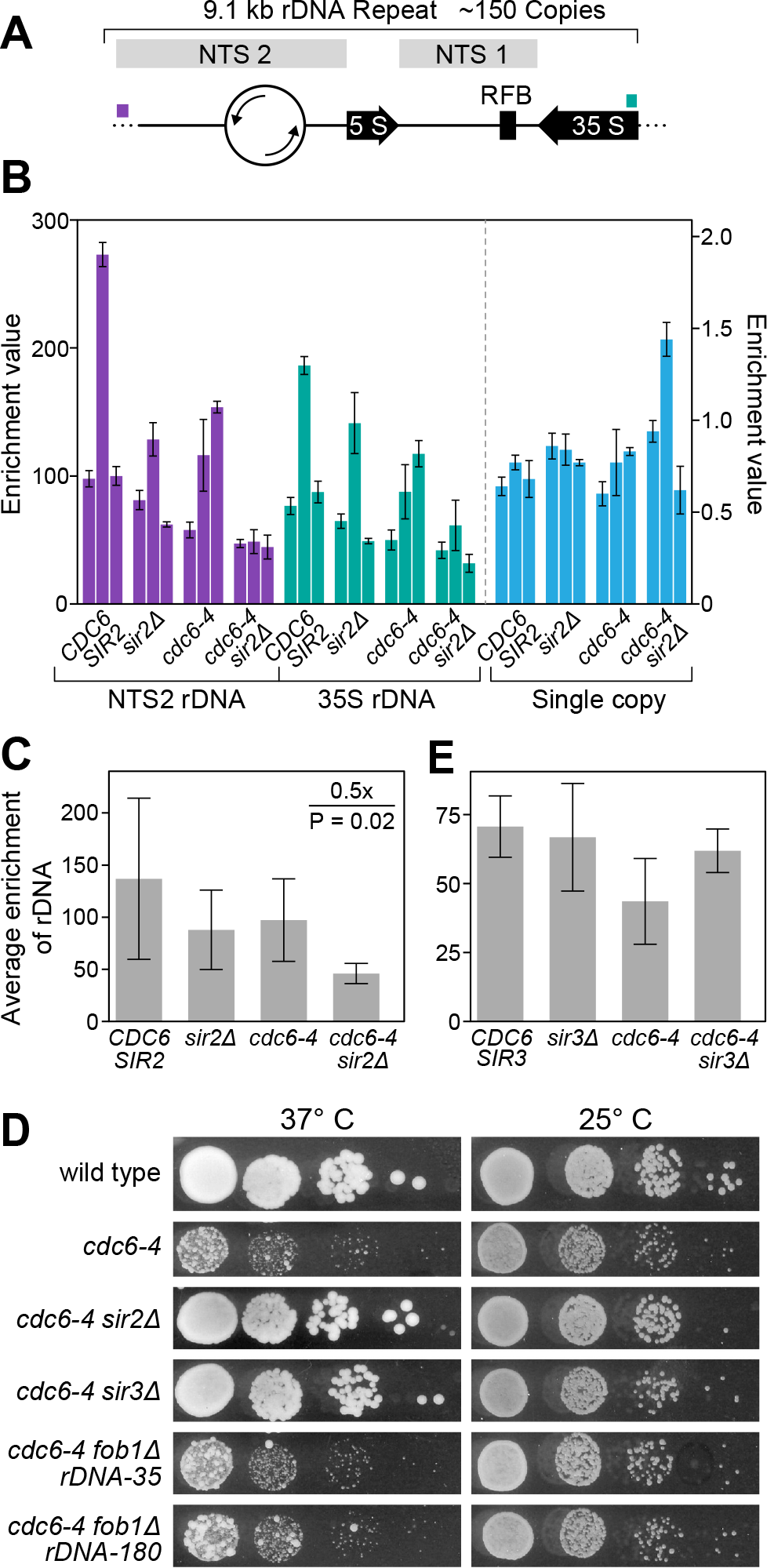
Reduction in rDNA copy number could not account for the suppression of *cdc6-4* by *sir2* mutants. **A.** Diagram of a single rDNA repeat is shown with the location of the primer pairs used to assess rDNA copy number by qPCR indicated by colored boxes. The rDNA origin is indicated as the open circle. NTS are the non-transcribed regions in the rDNA locus. RFB is the replication fork block element that binds Fob1. **B.** Enrichment values determined for the indicated strains and primer pairs. For each strain, three independent colonies were assessed (biological replicates), each represented by a bar with the standard error. Because we observed substantial variation in rDNA copy number between colonies from the same strains, we also used qPCR to assess the copy number of a single copy gene (*RIM15*) for the same DNA preparations (turquoise) to get some idea of the technical noise this assay produced. Note the different scale for the *RIM15* experiment as this value should theoretically equal 1.0. **C.** *SIR2* effect on the average enrichment values of the rDNA locus for each of the strains assessed in C. **D.** Growth of 10-fold dilutions of the indicated strains was assessed on solid YPD media at the indicated temperatures. **E.** The effect of *SIR3* on the average enrichment values of the rDNA locus for each of the indicated strains.

**Origin-specific depletion of acetylated H4K16-containing nucleosomes**. The pervasive yet origin-specific rescue of suppression of the MCM loading defect in *cdc6-4* cells by *sir2Δ* prompted us to consider the possibility that Sir2 might function directly on origin-adjacent nucleosomes within euchromatin. Sir2 is a deacetylase with specificity for acetylated lysine 16 of histone H4 (H4K16ac) [19]. Therefore, we used a comprehensive genome-wide histone modification atlas generated by high resolution MNase-ChIP-Seq of yeast nucleosomes to examine the acetylation status of nucleosomes adjacent to euchromatic origins [32] (**Figure 3**). To perform this analyses, we focused on two distinct groups of loci: experimentally-confirmed origins and non-origin intergenic regions that contain a match to the ORC binding site [43]. At confirmed origins, ORC and MCM binding, as well as origin activity, are experimentally documented, while at non-origin intergenic regions with ORC site matches, ORC and MCM binding are not detectable in vivo, and no origin function has been detected (**Figure 3A** non-origins n=179; origins; n = 259). 1201 bp fragments from these two distinct groups were aligned with the T-rich strand of the ORC site on the top strand in the 5’ to 3’ orientation (**Figure 3B**). *HM* silencer, telomeric (as defined by origins within 15 kb of chromosome ends) and rDNA origins were excluded from these and all subsequent analyses because these regions are known to be substantially enriched for H4K16 hypoacetylated nucleosomes as part of the *SIR*-dependent silencing mechanism. To confirm that we could use these data to recapitulate previously published results about nucleosome occupancy differences between origin and non-origin loci [43,44], nucleosome occupancy around the ORC site was assessed using the MNase-ChIP-Seq data generated from this more recent study [32]. The results from our analyses confirmed the conclusion that origin-adjacent nucleosomes show high occupancy at more defined positions around the nucleosome-depleted ORC site compared to the control non-origin adjacent nucleosomes (**Figure 3C**). Next, the H4K16ac status of nucleosomes surrounding the ORC site for both groups was determined and normalized to the H4K16ac status from a collection of nucleosomes present in a distinct collection of euchromatin intergenic regions that contained neither origins nor matches to the ORC site (n=239). This analysis revealed that nucleosomes adjacent to origins were depleted for H4K16ac but that nucleosomes adjacent to the control non-origin nucleosomes were not (**Figure 3D**). H3K9ac is also a potential substrate for Sir2 but, in contrast to a H4K16Q substitution, an H3K9Q substitution fails to suppress *cdc6-4* [19,23]. Also in contrast to H4K16ac, H3K9ac was depleted similarly from nucleosomes adjacent to origin and non-origin control loci (**Figure 3D**). Thus, H4K16ac was distinct among nucleosome acetylation marks in showing origin-specific depletion (**S3 Figure**).

**Figure 3.**
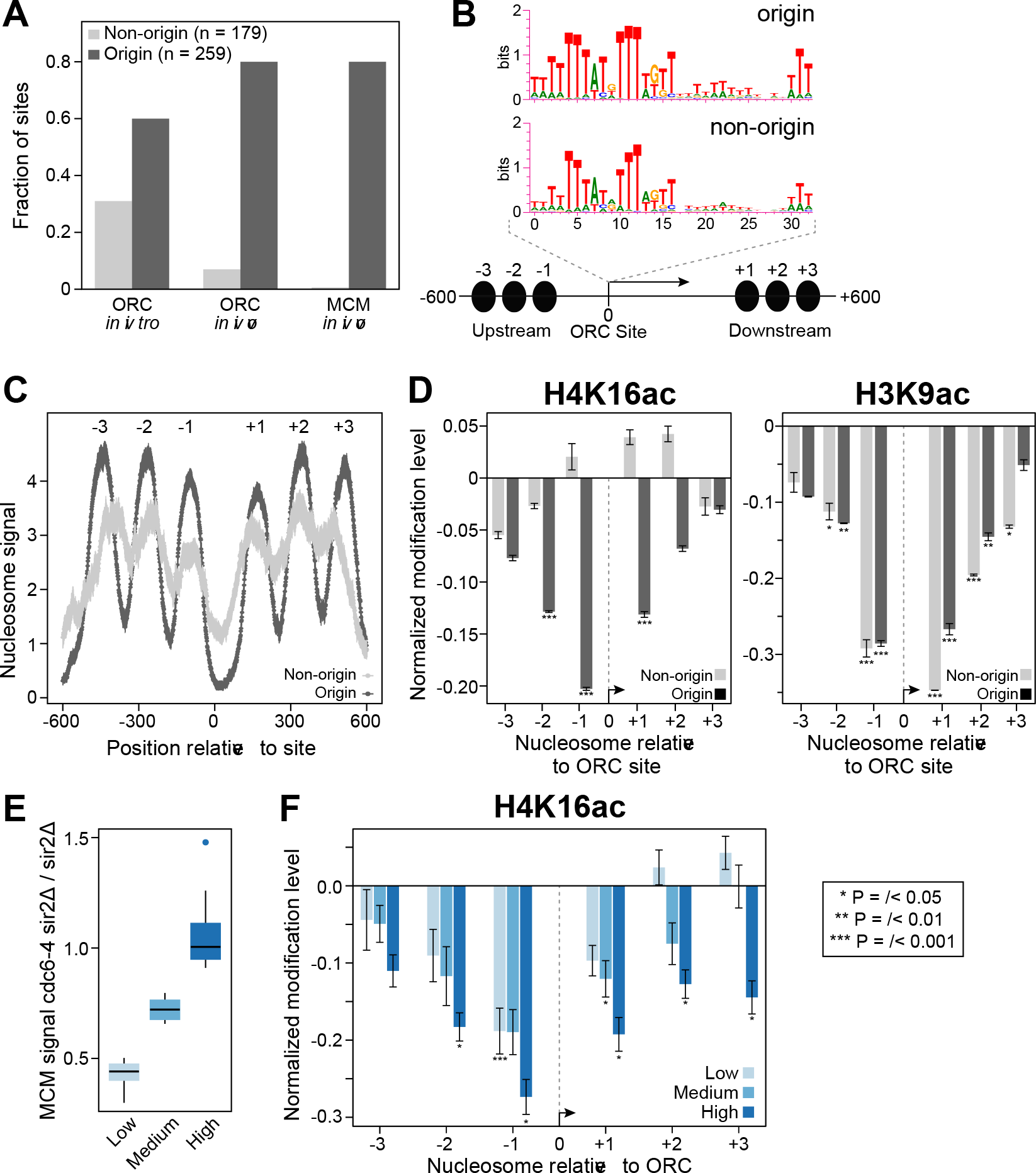
Acetylated H4K16 is depleted from origin-adjacent nucleosomes. **A.** Two groups of loci were analyzed, origins, defined as experimentally-confirmed origins with a confirmed or high-confidence ORC site, n = 259, and intergenic non-origin loci with ORC site matches but for which no origin activity has been observed, n = 179 [43]. “ORC in vitro” refers to loci that were bound by purified ORC in a genomic electrophoretic mobility assay [64]; “ORC in vivo” and “MCM in vivo” refer to the fraction of sites in these two groups that were detected by ORC- or MCM-ChIP, respectively. **B.** The WebLogo consensus for the ORC sites (or matches) in origins and non-origins, respectively, are shown above the diagram of the fragments used in the analyses of adjacent nucleosomes. Each fragment analyzed was oriented with the T-rich strand of the ORC site 5’ to 3’ on the top strand, and the first nucleotide of the ORC site was designated as position “0”. The fragments were 1201 bp such that six proximal nucleosomes, shown as black ovals, three on each side of the ORC site, were assessed. **C.** Nucleosome occupancy surrounding the origin and non-origin nucleosome depleted regions are shown using the MNase-ChIP-Seq nucleosome occupancy data from [32]. **D.** Normalized H4K16ac and H3K9ac for each of the six nucleosomes for the two different groups of loci examined. P-values are derived from Student’s T-test comparing the mean of acetylation status between each nucleosome to the mean acetylation status of nucleosomes from the 239 intergenic control loci 239 intergenic control loci. **E.** *SIR2*- responsiveness was defined as the ratio of the MCM ChIP-Seq signal in *cdc6-4 sir2Δ* to *sir2Δ* cells. The origins were ranked based on *SIR2*-responsiveness and then divided into quintiles, with the high quintile containing the most *SIR2*-responsive origins. **F.** H4K16ac status for the three quintiles of *SIR2*-responsive origins indicated in ‘**E**’ was determined as in ‘**D**’.

If depletion of H4K16ac on origin-adjacent nucleosomes was relevant to *SIR2*- dependent inhibition of MCM loading in *cdc6-4* cells, then the origins most responsive to deletion of *SIR2* might be expected to show the greatest depletion of H4K16ac. We defined origin *SIR2*-responsiveness as the ratio of the MCM ChIP-Seq signal in *cdc6-4 sir2Δ* cells to that in *sir2Δ* cells; the most *SIR2*-responsive origins generated ratios near 1.0, suggesting that deletion of *SIR2* substantially rescued the MCM loading defect of *cdc6-4* (**Figure 3E**). Comparison of H4K16ac status of nucleosomes surrounding the low, medium, and high-*SIR2* responsive origins revealed that as a group the most *SIR2*-responsive origins exhibited the greatest depletion of H416ac, especially when considering all six origin-adjacent nucleosomes (**Figure 3F**). Thus the varying degrees of responsiveness to *SIR2* for MCM association in *cdc6-4* cells correlated with the relative degree of H4K16 hypoacetylation at that origin.

**The depletion of acetylated H4K16-containing nucleosomes around euchromatic origins required *SIR2***. The data described above raised the possibility that Sir2 was deacetylating H4K16ac nucleosomes within euchromatin, specifically around origins. To test this possibility, we performed new H4K16ac MNase ChIP-Seq experiments on the exact same wild type (*SIR2*) and *sir2Δ* cells used for the MCM ChIP-Seq experiment described in Figure 1 (**Figure 4A**). Analyses of wild type cells recapitulated the published H4K16ac MNase ChIP-Seq results shown in Figure 3D, although we note the relative level of origin-specific H4K16ac depletion was slightly greater in these experiments, possibly due to differences in strain backgrounds or growth conditions. Regardless, the key result was that, in contrast to wild type, depletion of H4K16ac was lost from origin-adjacent nucleosomes in the *sir2Δ* cells, while the behavior of non-origin ORC site control nucleosomes was unchanged. Importantly, these effects also correlated with *SIR2* responsiveness (**Figure 4B**). Thus, origin-specific depletion of H4K16ac nucleosomes at euchromatic origins required *SIR2*.

**Sir2 and Sir3 were detected at origins**. Sir2 and Sir3 are physical components of yeast heterochromatin that have been detected at rDNA (Sir2) and telomeres and HM loci (Sir2 and Sir3) by ChIP experiments in multiple studies (reviewed in [15]). However, neither protein has been reported to associate with euchromatic origins [45]. The level of H4K16 hypoacetylation at euchromatic origins was estimated to be between 25-40% of that detected at heterochromatic origins for the +1 nucleosome (**S4 Figure**). This observation is consistent with a proteomic analysis of nucleosome modifications on a plasmid based origin [46]. However, given that Sir2 is an enzyme, stable binding by Sir2, or even Sir3 if it is required only transiently, might not be required to establish or maintain this modification state. To test this possibility, we used more recently published high-resolution ChIP-Seq Sir2 and Sir3 data sets and excluded all nucleosomes from known heterochromatin domains for normalization, as above, so that the baseline would not be affected by the extensive amount of Sir2 and Sir3 binding known to occur at these domains [34,47]. This analysis detected Sir2 and Sir3 ChIP-Seq signals on nucleosomes adjacent to origins but not to non-origin controls (**Figure 5A**). Because the Sir3 data was generated from a high-resolution MNase ChIP-Seq experiment, we also examined these data at nucleotide resolution across the 1201 bp origin and non-origin fragments described in Figure 3B, normalizing the number of reads that contained a given nucleotide in the ChIP sample to the total number of reads that contained that nucleotide in the starting material, as in [48] (**Figure 5B**). These analyses revealed a Sir3 ChIP-Seq signal that was strongest at the most proximal origin-adjacent nucleosomes (−1 and +1) but also detectable above baseline at the −3, −2, +2 and +3 nucleosomes (**Figure 5C**). As was true for the depletion of H4K16ac, the Sir3 signals correlated with the degree of *SIR2*-responsiveness as defined in Figure 3E (**Figure 5D-E**). Randomizing the origins into three different sets prior to plotting the Sir3 ChIP-Seq signals eliminated the correlation (**Figure 5F**). Taken together with the data presented in Figures 3 and 4, several molecular hallmarks of heterochomatin – Sir2, Sir3 and Sir2-dependent H4K16 hypoacetylation – could be detected at euchromatin origins but not at euchromatic non-origin controls. In addition, the relative levels of each of these molecular hallmarks correlated with how strongly a *SIR2* deletion restored an MCM ChIP signal to origins in *cdc6-4* cells grown at temperatures that abolished MCM loading.

**Figure 4.**
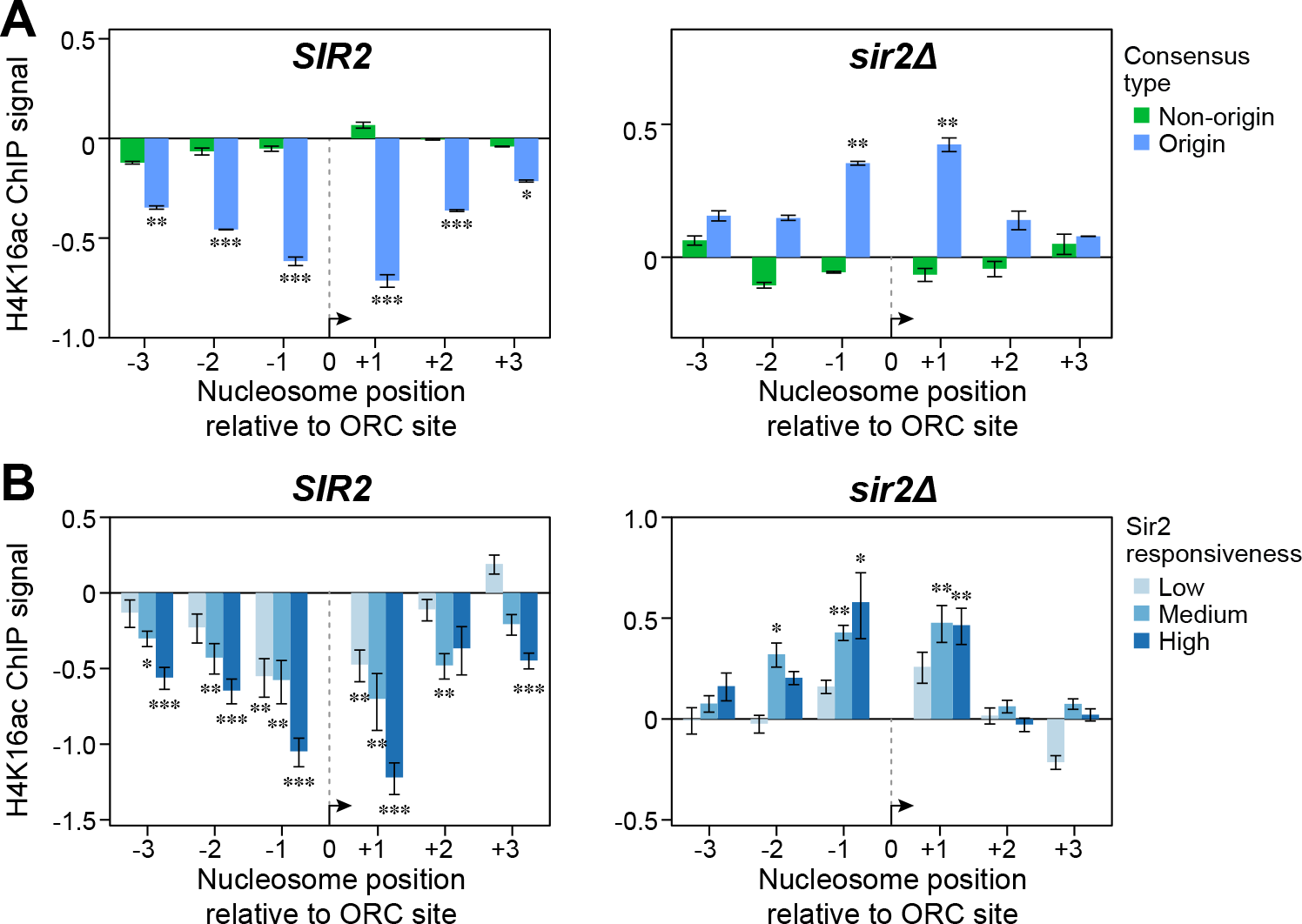
Depletion of H4K16ac from origin-adjacent nucleosomes required *SIR2*. H4K16 acetylation status of nucleosomes was assessed by MNase ChIP-Seq in **A.** *SIR2* and *sir2Δ* cells used in Figure 1. These cells were the same as those analyzed for MCM binding in Figure 1. **B.** The H4K16 acetylation status was plotted for the low, medium and high SIR2-responsive quintiles as defined in Figure 3.

**Figure 5.**
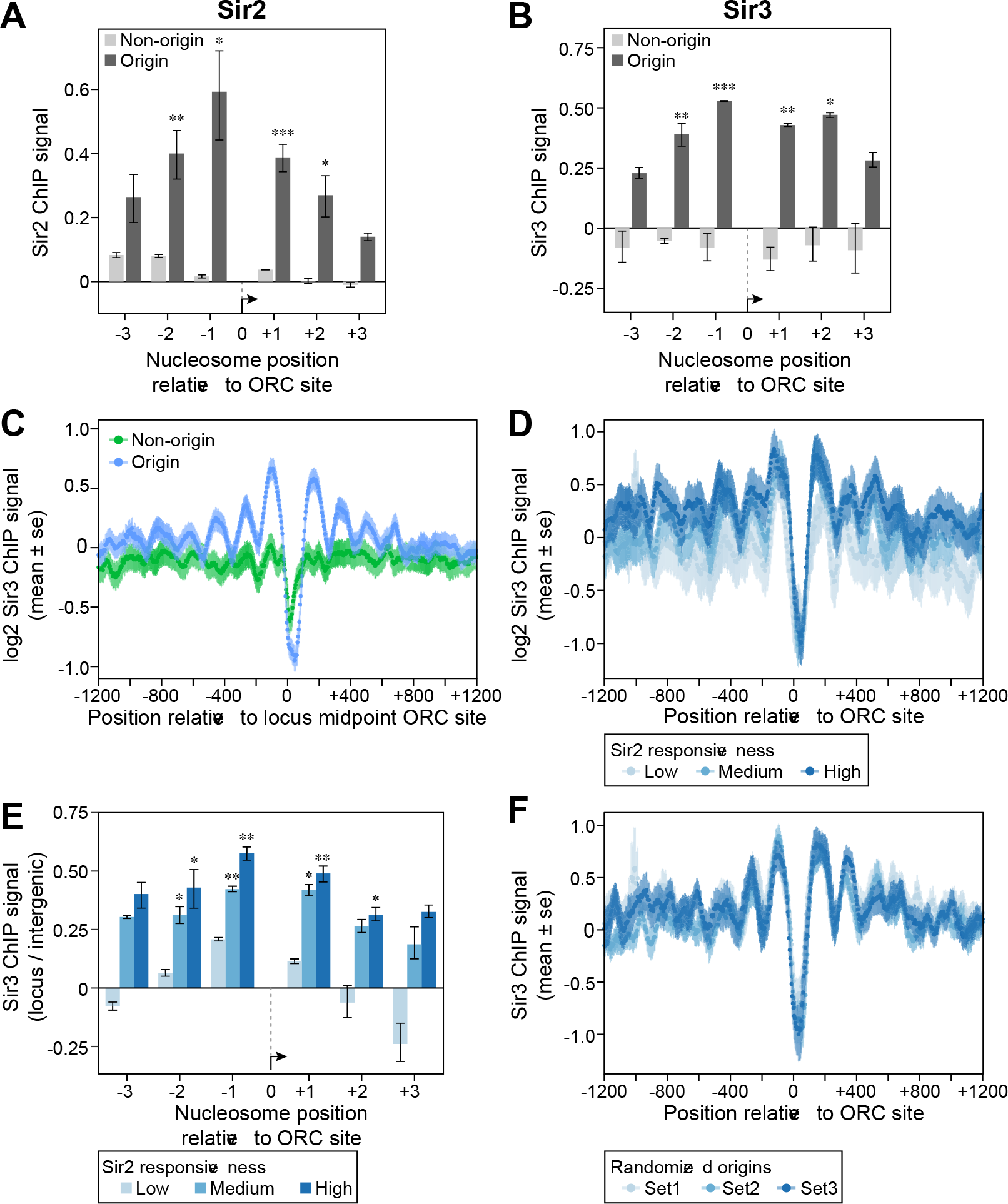
Sir2 and Sir3 were physically associated with nucleosomes adjacent to origins but non-origin controls. **A.** Sir2 or (**B)** Sir3 binding to origin or non-origin adjacent nucleosomes was assessed as in Figure 3 using data from [33,34]. **C.** The normalized Sir3 ChIP-Seq signal (log_2_ Sir3 ChIP signal, y-axis) for each nucleotide was plotted over the 1201 bp origin (blue) or non-origin (green) loci (coordinates on x-axis). The Sir3 ChIP-Seq signal represented total reads for each nucleotide normalized to the depth (total reads) and breadth (number of nucleotides with reads) of the sequencing reactions [48]. Genomic regions previously established as Sir2-heterochromatic domains were excluded from the normalization. **D. and E.** The Sir3 ChIP-Signals were plotted as in ‘**C**’ or ‘**B**’, respectively, for the three different *SIR2*-responsive groups defined in Figure 3. **F.** The *SIR2*-responsive origins comprising the three groups assessed in ‘**D’** were randomized into three different sets and the Sir3 ChIP-Seq signals determined and plotted as in ‘**D**’.

**A *cdc6-4* suppressor screen using core modifiable histone H3 and H4 mutants identified residues important for Sir3 binding**. Histones H3 and H4 form a tetramer that binds both to dsDNA and to two dimers of histones H2A and H2B to form the nucleosome. Previous work showed that many conserved residues on the histone H3 and H4 N-terminal tails as well as within the globular core region (“core-modifiable” residues) can be post-translationally modified. As discussed above, an H4K16Q substitution that should mimic the acetylated form of this residue (i.e. the form that could theoretically mimic the effect of a *sir2Δ*), suppressed the temperature-sensitive growth defect of *cdc6-4* [23]. To test whether we could identify additional alleles with similar suppressive behavior, we assessed a library of 46 histone H3 and H4 mutations affecting 16 distinct core-modifiable residues for suppression of the *cdc6-4* growth defect (**Figure 6A)**. Mutant plasmids harboring a single copy of the *HHT2-HHF2* locus marked with *TRP1* were transformed into wild type and *cdc6-4* cells lacking both chromosomal copies of histone H3 and H4 genes, maintained by a *HHT2-HHF2 URA3 ARS CEN* plasmid, and colonies were subsequently plated on 5-FOA containing media to select against the wild type *HHT2-HHF2* plasmid. Recovered cells were then examined for growth on YPD medium at 25°C and higher temperatures. In agreement with previous studies, none of the histone mutants, with the exception of H4-S47E, caused growth defects in wild type cells [49], and several mutants were identified that inhibited growth of cells harboring the *cdc6-4* allele even at the permissive growth temperature (25°C), (SS and MW, unpublished). However, relevant to this study, specific substitutions at only two residues, H3K79 (H3K79A and H3K79E but not H3K79R) and H4K79 (H4K79A and H4K79R) suppressed the temperature-sensitive growth defect of *cdc6-4* cells (**Figure 6B**). Notably, these residues are important for Sir3 binding to the nucleosome and for Sir3-mediated transcriptional silencing at the *HM* and telomeric loci [50–52]. H3K79 is a substrate for methylation by Dot1 and Sir3 binds preferentially to unmethylated H3K79 [53,54]. Alanine mutations at 4 residues adjacent to H3K79 (H3E73A, H3I74A, H3T80A and H3D81A) and a charge reversal mutation (H3E73K) were also screened. This revealed that mutations at H3E73 and H3T80 also suppressed the temperature-sensitive growth defect of *cdc6-4* cells (**Figure 6B**). Mutations at H3E73 and H3T80 were previously shown to significantly disrupt silencing at the *HM* and telomeric loci [50] and notably, H3T80 directly contacts the LRS domain of Sir3 but H3D81 does not [51,52]. The four residues whose substitutions suppressed *cdc6-4* ts lethality (H3E73, H3K79, H3T80, and H4K79) clustered to a small patch on the nucleosome surface critical for binding Sir3 directly (**Figure 6C**, **highlighted in red**) [51]. Thus disruption of Sir3 binding to nucleosomes suppressed the *cdc6-4* growth defect. Together with the ChIP-Seq data, these genetic data supported the conclusion that both Sir2 and Sir3 directly alter the molecular features of nucleosomes adjacent to euchromatic origins, thus generating a local chromatin environment that can negatively impact the MCM loading reaction.

**Figure 6.**
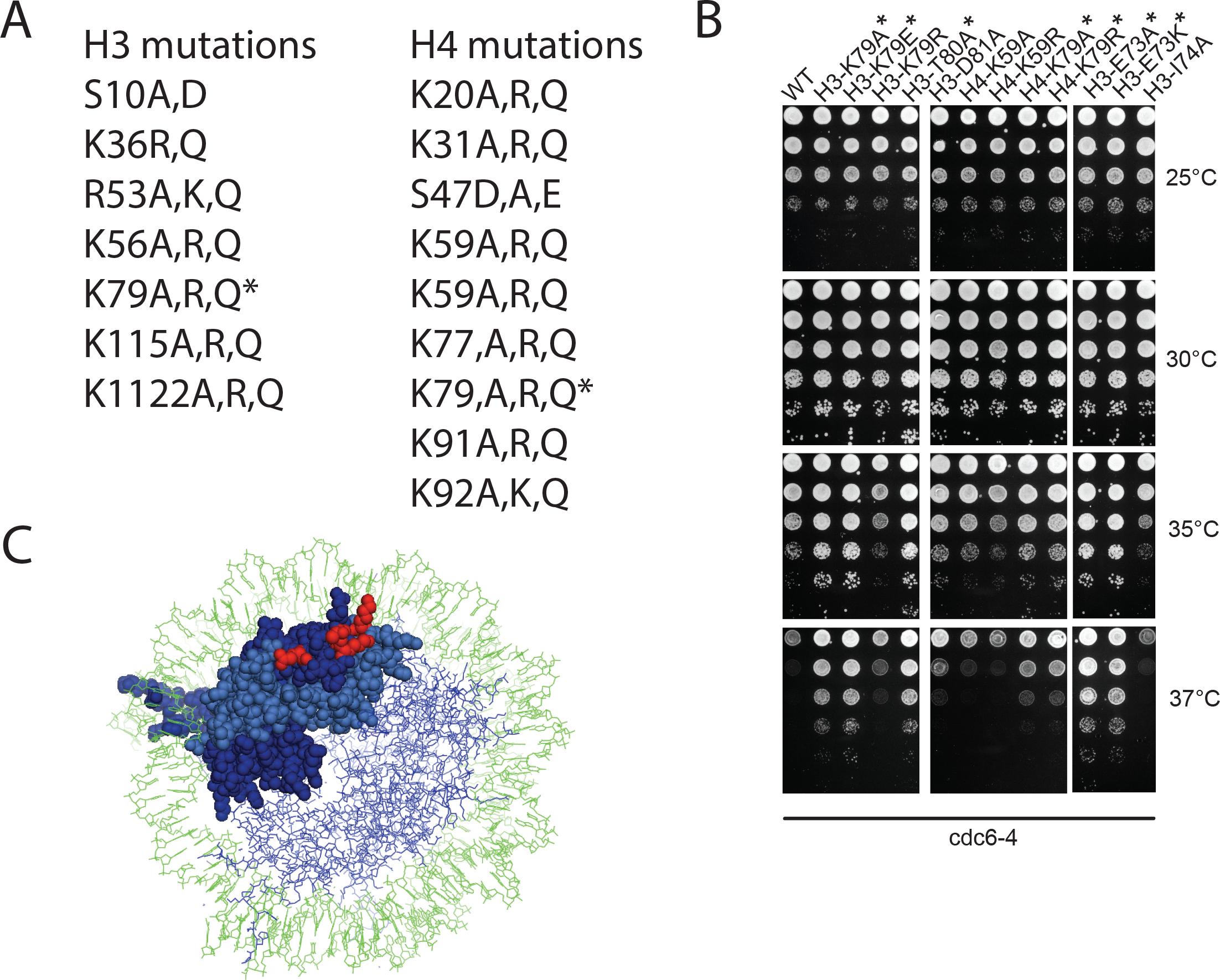
Histone H3 and H4 core modifiable alleles that suppress *cdc6-4* temperature sensitivity define a known Sir3-nucleosome interface. **A.** List of histone point mutations that were initially screened for suppression of *cdc6-4* temperature sensitivity. Asterisks indicate the only residues when mutated that gave suppression. **B.** 10-fold serial dilutions of M1345 (*cdc6-4*) with either wild type H3, H4 or various histone mutants, were spotted onto YPD and grown at the indicated temperatures. H4K59A and H4K59R as shown as examples of histone mutants that did not suppress. **C.** Nucleosome structure (PDB-1ID3) with one H3(dark blue)-H4(light blue) dimer highlighted with the H3E73, H3K79,

## Discussion

Sir2 and Sir3 are non-essential proteins known best for their direct functions in heterochromatin-mediated transcriptional gene silencing (reviewed in [15]). This report described a new, pervasive and direct heterochromatin-independent role for these regulators at euchromatic DNA replication origins. The role was pervasive in that Sir2 exerted a negative effect on the MCM loading reaction at most origins in the genome. The evidence in support of this conclusion came from a comparison of MCM loading at origins by ChIP-Seq in *cdc6-4* mutant cells and *cdc6-4 sir2Δ* cells. Cdc6 is essential for the MCM complex loading reaction, and the *cdc6-4* allele weakens Cdc6 function such that, under non-permissive growth temperatures, detectable MCM association was abolished from all origins. However, under the same conditions, *cdc6-4 sir2Δ* cells exhibited MCM association at most euchromatic origins (~80%). Thus, the ability of *sir2* inactivating mutations to robustly suppress the temperature-sensitive lethality of *cdc6-4*, as well as growth defects caused by other replication alleles that reduce or abolish MCM loading [29], is due to genome-scale enhancement of MCM loading at most euchromatic origins.

The ability of *sir2Δ* to enhance MCM complex loading at most euchromatic origins in *cdc6-4* cells was striking, but it did not indicate whether this effect was due to Sir2 acting directly within euchromatin in general or at euchromatic origins in particular. Indeed, recent reports establish that *SIR2* can alter origin function within euchromatin in yeast indirectly because of its function in rDNA heterochromatin formation that suppresses many of the rDNA repeat origins [22,25]. However, analyses of the rDNA locus presented here suggested that a similar rDNA-mediated mechanism could not explain why *sir2Δ* or *sir3Δ* suppressed *cdc6-4* so robustly. Instead, the data provided compelling evidence that Sir2 functioned directly at euchromatic origins, and that it was this direct function of Sir2 that made yeast so vulnerable to defects in the MCM complex loading reaction caused by *cdc6-4*. First, relative hypoacetylation of H4K16 was observed for nucleosomes immediately adjacent to euchromatic origins but not for nucleosomes adjacent to non-origin control loci. Importantly, these nucleosome states were observed in wild type cells (i.e. not the *cdc6-4* mutant). In addition, of the twelve histone acetylation marks examined [32], H4K16ac was the only one that showed this origin-specific depletion. Second, origin-adjacent nucleosomal H4K16 hypoacetylation required *SIR2*; a *sir2Δ* mutant completely lost the nucleosomal H4K16 hypoacetylation pattern at euchromatic origins, without having any obvious effect on nucleosomes adjacent to the non-origin control loci. Third, analyses of previously published high-resolution ChIP-Seq experiments [34,55] using the same computational approach that was used to examine H4K16ac, provided evidence that Sir2 and Sir3 were also physically associated with nucleosomes adjacent to euchromatic origins but not non-origin control loci. Sir2 and Sir3 exhibited the same pattern but in the exact opposite direction of H4K16 acetylation, which is what is observed for these proteins and this histone mark at silenced domains. Lastly, each of these defining molecular features of classic SIR-heterochromatin correlated with *SIR2*-responsiveness – defined as the relative extent of MCM association rescue at origins in *cdc6-4 sir2Δ* cells. To the best of our knowledge, these data provide the first evidence that the heterochromatin proteins Sir2 and Sir3 act directly on the local chromatin environment of euchromatic origins, and therefore define a new role for these proteins.

Despite the common molecular hallmarks shared between the Sir2-3-chromatin at euchromatic origins described in this study and classic *SIR*-mediated heterochromatin, these two types of chromatin domains are different. For example, unlike *SIR*- heterochromatin that functions in gene silencing at the *HM* loci, telomeres or rDNA, there is no evidence that origin-adjacent chromatin functions as a robust transcriptionally silenced domain. In addition, *SIR*-heterochromatin affects multiple nucleosomes, defining domains that encompass ~4-10kb of contiguous chromosomal DNA (reviewed in [15]), whereas the Sir2-3-chromatin at origins encompassed <1 kb of chromosomal DNA adjacent to the origin and affected six nucleosomes at most. While it remains to be determined whether the specific molecular interactions that recruit Sir2 and Sir3 to euchromatic origins are also used to recruit these proteins to heterochromatic regions, it is clear that their functional and structural impact on nucleosomes at euchromatic origins is attenuated relative to their impact within heterochromatin. We suggest that the Sir2-3 chromatin at origins defines a silencing-independent role for these proteins.

**A specific G1-phase role for H4K16 acetylation at origins**: Acetylation of histone H3 and H4 N-terminii at origin proximal-nucleosomes generally enhances origin function [20,56,57]. However, it has been difficult to assign specific roles of this acetylation to individual histone H3 or H4 lysines. Indeed, in terms of origin activation, while acetylation of histone H3 and H4 tail lysines is clearly important for origin activation during S-phase, no single lysine acetylation event is sufficient [46]. In addition, several different combinations of multiple lysine to arginine substitutions on the histone H3 and H4 N-terminal tails can substantially reduce origin activation, suggesting that some threshold level of nucleosome acetylation, or the concomitant charge neutralization, is what is important for origin activation. In contrast, the initial identification of a *sir2* mutation, and subsequently a H4K16Q substitution, as robust suppressors of *cdc6-4* temperature-sensitivity and origin-specific MCM loading, indicated that H4K16 acetylation might be unique among histone tail lysines in being particularly relevant to origin licensing [23,29]. This study revealed that H4K16 was also unique among histone H3 and H4 tail lysines because H4K16 exhibited *SIR2*-dependent, origin-specific hypoacetylation on nucleosomes immediately flanking euchromatic origins. Consistent with these observations, proteomic analysis of nucleosomes adjacent to a plasmid-origin reveals that H4K16 behaves uniquely among histone H3 and H4 tail acetylation marks [46]. Specifically, in G1-phase, nucleosomes adjacent to a plasmid-based origin were relatively hypoacetylated at H4K16 compared to bulk nucleosomes, whereas the other histone tail lysines analyzed (H3-K9, −K14, K-23; H4-K5, −K8, −K12) showed similar levels of hypoacetylation on bulk and plasmid-based origin adjacent nucleosomes. Because Sir2 and Sir3 also showed origin-specific association with nucleosomes, the distinctive role for H4K16 acetylation in control of the G1-phase MCM loading reaction can now be explained by formation of a Sir2-3 chromatin structure directly at many euchromatic origins, suggesting that origins, perhaps via ORC, can specifically recruit Sir2-3.

**MCM loading challenges in a native chromatin context**: The data presented here provided evidence that the MCM loading reaction has evolved to work within a naturally repressive chromatin environment established by Sir2-3. Indeed, an otherwise severely crippled licensing reaction caused by the *cdc6-4* defect can quite effectively license most origins if the native repressive chromatin environment is abolished by inactivation of Sir2. While the precise mechanistic step(s) of the MCM complex loading reaction that are affected by the Sir2,3-repressive chromatin environment remain to be examined, recent successes in reconstituting DNA replication in chromatin contexts in vitro offer an ideal pathway forward [58–60]. For example, while reconstituted chromatin does not block MCM complex loading in vitro, it would be interesting to learn whether the addition of Sir3 to these biochemical reactions is sufficient for inhibition and, if so, requires Sir3’s ability to bind to nucleosomes [51].

However, in terms of yeast physiology, the major challenge moving forward is to understand why such a chromatin-structure capable of inhibiting MCM complex loading may have evolved to exist at so many euchromatic origins at all. While a *sir2Δ* or a *sir3Δ* have no obvious genome-scale effect on yeast cell division, MCM complex loading or origin activation in cells with wild type MCM complex loading reaction components, the Sir2,3-chromatin structure discussed here does indeed exist in wild type cells, and, based on the strong genetic suppression of *cdc6-4* and other alleles that weaken MCM complex loading [29], is clearly capable of exerting a substantial inhibitory effect on this reaction. Indeed, the results presented here mean that the levels and or activities of the MCM complex loading proteins have evolved to contend with a particularly inhibitory local chromatin structure around euchromatic origins; a temperature-sensitive allele that weakened MCM complex loading, such as *cdc6-4*, would never have been isolated in yeast cells lacking Sir2 or Sir3 as the mutant Cdc6-4 protein provides for robust function in such contexts. It is possible that the Sir2,3- chromatin at euchromatic origins is simply a byproduct of yeast cells evolving transcriptionally silenced heterochromatic domains in the first place, and, as a result of strong evolutionary selection for these regions’ existence and the fundamental biochemistry of protein dynamics, Sir2 and Sir3 simply fall off of heterochromatin at some frequency and, when they do, end up concentrating at origins for a time as they bounce around the nucleus [61–63]. Because of this unavoidable biochemical reality, the MCM complex loading reaction had to evolve more robustly than it would have otherwise. Additionally, and/or alternatively, the Sir2,3- chromatin at euchromatic origins may play an important role in normal yeast physiology, offering a fine-tuning of the MCM complex loading reaction at many individual origins that, while at the first-level of cell growth and origin function in bulk laboratory assays may be hard to measure, nevertheless has important repercussions at the population and/or evolutionary scale for yeast and other eukaryotic organisms. In this regard, we note that human SIRT1, the ortholog of yeast Sir2, can suppress the function of dormant origins under conditions of replication stress, suggesting an evolutionary conserved, and thus physiologically relevant role for heterochromatin regulatory proteins at euchromatic origins. The increasing availability of higher-resolution experiments will allow for more stringent examination of the role of Sir2,3-chromatin at so many euchromatic origins in yeast physiology.

## Material & Methods

**Yeast strains**. The strains used for plasmid stability measurements and Mcm2 ChIP-Seq were the W303-1A derivatives: M138 (*MAT***a**), M386 (*MAT***a** *cdc6-4::LEU2*), M922 (*MAT***a** *cdc6-4::LEU2 sir2Δ::TRP1*) (described in Pappas et al. 2004), M1014 (*MAT***a**
*cdc6-4*) and M2126 (*MAT***a** *sir2Δ::TRP1*). The M1321 (WT) and M1345 (*cdc6-4*) histone shuffle strains were described previously (Crampton et al., 2008). The strains used for genetic suppression and/or rDNA copy number experiments (Figure 2) include, in addition to M138, M386, M922 and M2126 described above, CFY43 (*MAT***a** *FOB1 CDC6 SIR2 sir3F::TRP1*), CFY4584 (*MAT***a** *fob1 Δ::HIS3 CDC6 SIR2 SIR3 rDNA-35*), CFY4585 (*MAT***a** *fob1 Δ::HIS3 CDC6 SIR2 SIR3 rDNA-180*), CFY4603 (*MAT***a** *fob1 Δ::HIS3 cdc6-4 SIR2 SIR3 rDNA-35*), CFY4604 (*MAT***a** *fob1 Δ::HIS3 cdc6-4 SIR2 SIR3 rDNA-180*), CFY4613 (*MAT***a** *FOB1 cdc6-4 SIR2 sir3Δ::TRP1*. CFY4583 and CFY4585 were provided by Jonathan Houseley (Babraham Institute, Cambridge, UK) [40].

**ChIP-Seq Experiments**: For the MCM ChIP-Seq experiments, yeast cells were grown in liquid YPD at 25°C from single colonies until they reached an A600 of 0.2, at which point nocodazole was added to a final concentration of 15ug/ml and the cultures incubated for 2.5 hours, to obtain a uniform G2/M-phase arrest. Cultures were shifted to 37°C for 30 minutes, the non-permissive growth temperature for *cdc6-4*, and then released from the nocodazole arrest at 37°C. When the majority of cells had entered G1-phase (55- minutes post-release for *CDC6 SIR2*, *sir2Δ* and *cdc6-4 sir2Δ* and 110 minutes for *cdc6-4*), OD equivalents of each cell line were cross-linked with formaldehyde for 15 minutes and then harvested, with chromatin prepared for IP, by sonication for ChIP-Seq. Mcm2 ChIP was performed using a monoclonal antibody raised against yeast Mcm2 (gift of Bruce Stillman, Cold Spring Harbor Laboratory). Three independent biological replicates and two technical replicates for each biological replicate were performed for each strain. The ChIP DNA was prepared for deep sequencing by the UW-Madison Biotechnology Center using http://www.biooscientific.com/Portals/0/Manuals/NGS/5143-01-NEXTflex-ChIP-Seq-Kit.pdf. Sequencing was done on the HiSeq2000 1×100. Quality and quantity of the finished libraries were assessed using an Agilent DNA1000 chip and <u>Qubit^®^ dsDNA HS Assay Kit</u>, respectively. Libraries were standardized to 2nM. Cluster generation was performed using standard Cluster Kits and the Illumina Cluster Station. Single-end, 100bp sequencing was performed, using standard SBS chemistry on an Illumina HiSeq2000 sequencer. Images were analyzed using the standard Illumina Pipeline, version 1.8.2. Sequencing data for this project are available at the NCBI BioProject ID PRJNA428768.

For the downstream analyses (**Figure 1** **and S1**), all data for a given strain were combined as each replicate generated virtually identical reads. However, for the *cdc6-4* strain, which produced extremely low signal-to-noise data, only two technical replicates from a single biological sample were used. For downstream analyses using MochiView the CG1 format data was collated into 25 bp bins (**Figure 1**). The combined data from each strain was run through CisGenome to identify the top 1000 peaks, divided into 100 peak bins and the percent of peaks that overlapped with at least one ARS (ARS coordinates used from oriDB) determined. The oriDB lists 410 yeast origins as confirmed. Based on these data and the above analyses we chose to examine and compare the top 400 peaks from each sample for our initial analyses (**Figure S1**). The raw data for *CDC6 SIR2*, *sir2Δ* and *cdc6-4 sir2Δ* were normalized and scaled to the same *cdc6-4* data for to generate the final MCM signal values for downstream analyses.

The H4K16-acetylation MNase-ChIP DNA was generated from yeast strains M138 (*CDC6 SIR2* strain used in Figure 1) and M2126 (*CDC6 sir2Δ* strain used in Figure 1) following the protocol described in [32], except a Bio101 Thermo FastPrep FP120 was used to break cells after crosslinking. Purified DNA was submitted to the University of Wisconsin-Madison Biotechnology Center. DNA concentration and sizing were verified using the Qubit^®^ dsDNA HS Assay Kit (Invitrogen, Carlsbad, California, USA) and Agilent DNA HS chip (Agilent Technologies, Inc., Santa Clara, CA, USA), respectively. Samples were prepared according the TruSeq^®^ ChIP Sample Preparation kit (Illumina Inc., San Diego, California, USA) with minor modifications. Libraries were size selected for an average insert size of 350 bp using SPRI-based bead selection. Quality and quantity of the finished libraries were assessed using an Agilent DNA1000 chip and Qubit^®^ dsDNA HS Assay Kit, respectively. Libraries were standardized to 2nM. Paired end 300bp sequencing was performed, using SBS chemistry (v3) on an Illumina MiSeq sequencer. Images were analyzed using the standard Illumina Pipeline, version 1.8.2. Sequencing data for this project are available at the NCBI BioProject ID PRJNA428768. For the data shown in **Figure 4,** downstream analyses were performed as described (Weiner et al., 2015).

**rDNA copy number determination**: Quantitative PCR reactions were carried out in sealed 200 ul microplates in a BioRad C1000 Thermocycler, CFX96 Real-Time System. The conditions were: 95°C-3’|95°C −30’’, 58°C −15’’|Repeat 40 cycles|Standard Melt Curve Analysis. Each reaction contained: 0.04 ng/uL genomic DNA template, 0.016 nM of each primer, and 1X qPCR master mix (0.1 mM dNTPs, 1.25% fromamide, 0.1 mg/mL BSA, 0.5 U Taq DNA polymerase, 5 mM Tris-pH 8.0, 10 mM KCl, 1.5 mM MgCl2, 0.75% Triton and 0.5X SYBR Green DNA stain.) DNA concentration and primer efficiency validation were performed by testing primer efficiency by titrating template concentrations. Serial dilutions of DNA were used to test a range of DNA concentrations from 0.1 ng/uL to 0.0016 ng/uL in technical triplicates. Efficiency values for RIM15, NTS2 and NL primer sets were calculated as 83%, 83%, and 81%, respectively, by the BioRad CFX96 software. The entire range of DNA concentrations was found to be within the linear range of instrument response. Melt curve analysis revealed a single product from each primer pair that was confirmed in each experimental run. The primer pairs were: NTS2: GGGCGATAATGACGGGAAGA-Fwd and TGTCCACTTTCAACCGTCCC-Rev; 35S (D1/D2 loop of 26S rDNA): GCATATCAATAAGCGGAGGAAAAG-Fwd and ACTTTACAAAGAACCGCACTCC-Rev; RIM15: GCCAGAACATTGGGTCAGAT-Fwd and CCGGATACTCGGATGTGTCT-Rev.

**Histone H3 and H4 suppressor screen**. Strains M1321 (MATa ade2 ura3 leu2-3,112 trp1 hht1,hhf1::LEU2 hht2,hhf2::HIS3/pDP378 (HHT2,HHF2 CEN6 ARSH4 URA3) and M1345 (M1321, cdc6::ura3 LEU2::cdc6-4) were transformed with the *TRP1* histone H3, H4 plasmids pH3H4-WT (J. Boeke) and 26 mutant derivatives as shown in Figure 6A at 25°C. Multiple transformants were streak purified on FOA medium, recovered to YPD at 25°C and then screened for growth at 35°C and 37°C on YPD medium. Histone mutations were also constructed in pDP373 (pRS414 (TRP1) containing HHT2-HHF2 on a Spe1 fragment) using QuikChange. The entire H3 and H4 genes were sequenced to verify wild type sequences other than point mutation introduced, then transformed in M1321 and M1345 and used for spotting experiments in Figure 6.

## Acknowledgements

We are grateful to Bruce Stillman (Cold Spring Harbor Laboratory) for sharing Mcm2 antibodies and for critical discussions. We also thank the following people for discussions and critical advice: Elizabeth Kwan and M.K. Raghuraman (University of Washington, Seattle); Tsung-Han Hsieh and Oliver Rando (University of Massachusetts Medical School, Worcester); Xiaolan Zhao (Memorial Sloan Kettering Cancer Center, New York). Finally, we thank Jonathan Houseley (Babraham Institute, Cambridge, UK) for yeast strains, Christopher Warren (Proteovista LLC, Madison, WI) for the initial processing of the MCM ChIP-Seq data, and Josh Hyman and the Biotechnology Center, University of Wisconsin, Madison for sequencing and guidance. Support for this project was provided by NIH GM056890 to CAF, the Integrated Program in Biochemistry (University of Wisconsin, Madison, WI) (KP and JC), the Department of Biomolecular Chemistry (School of Medicine and Public Health, UW Madison) (TH and CAF) and by the Van Andel Institute and NSF MCB-0950464 to MW.

## Author Contributions

Design and Direction of Study: TH, CAF, MW; Contributions to Experimental Design and/or Execution: TH, FC, KP, SS, JK, JC, ES, JB, CAF, MW; Data Analysis: TH, CAF, MW; Writing and Editing of Manuscript: CAF, ES, TH, MW

